# Exploring of bacterial blight resistance in landraces, mining of resistant gene(s) and genetic divergence of resistant germplasm through molecular markers and quantitative traits

**DOI:** 10.1101/2021.08.04.455106

**Authors:** Touhidur Rahman Anik, Sheikh Arafat Islam Nihad, Md. Al-Imran Hasan, Mohammad Ahasan Hossain, Md. Mamunur Rashid, Mohammad Ashik Iqbal Khan, Mohammad Abdul Latif

## Abstract

Bacterial blight, one of the oldest and severe diseases of rice poses a major threat towards global rice production and food security. Thereafter, sustainable management of this disease has given paramount importance globally. In the current study, we explored 792 landraces to evaluate their disease reaction status against three highly virulent strains (*BXo69*, *BXo87* and *BXo93*) of *Xanthomonas oryzae* pv*. oryzae* (*Xoo*). Subsequently, we intended to identify the possible candidate resistant (*R*) genes responsible for the resistant reaction using six STS markers correspond to *Xa4*, *xa5*, *Xa7*, *xa13*, *Xa21* and *Xa23* genes and finally, we evaluated morphological and molecular diversity of the potential bacterial blight resistant germplasm using quantitative traits and ISSR markers. Based on pathogenicity test, a single germplasm was found as highly resistant while, 33 germplasm were resistant and 40 were moderately resistant. Further molecular study on these 74 germplasm divulged that 41 germplasm carried *Xa4* gene, 15 carried *xa5* gene, 62 carried *Xa7* gene, 33 carried *xa13* gene, and 19 carried *Xa23* gene. Only a single germplasm consisted of *Xa21* gene. Interestingly, we found a wide range of gene combinations ranged from 2 to 4 genes among the germplasm resistant to bacterial blight and G3 genotype (Acc. No. 4216; highly resistant) having *Xa4*, *Xa7*, *xa13*, *Xa21* and G43 genotype (Acc. No. 1523; resistant) having *Xa4*, *xa5*, *xa13* and *Xa23* gene combination being the most effective against all the *Xoo* strains. Nonetheless, UPGMA dendrogram and heatmap analysis based on quantitative traits identified two important clusters viz. cluster-III and cluster-VIII with multiple desired traits However, genetic similarity based on ISSR marker data pointed out 3 germplasm, namely, G20 (Acc. No. 4004; contained Xa4, Xa7, xa13 genes), G17 (Acc. No. 3981; contained Xa7, xa13 genes) and G6 (Acc. No. 991; contained Xa4, Xa7, xa13, Xa23 genes) having comparatively lower genetic similarity (0.30, 0.37 and 0.38, respectively) with elite variety BRRI dhan28. Notably, Mantle test of molecular and morphological data indicated that there was a positive correlation (r = 0.113) between them. The outcome of this study would enrich and diversify the rice gene pool and would be useful for the development of durable bacterial blight resistant varieties.

## Introduction

Rice, a monocot cereal grain, is considered as primary energy source for more than 3.9 billion people around the world. Annually more than hundred countries across the globe produce over 755 million tons of rice in roughly 162 million hectares of land [1]. Whereas Bangladesh, the fourth largest rice producing country annually produces about 54.64 million tons of rice, feeding its 135 million people whose half of total protein intake and two-third of overall calorie input come from rice [2]. However, worldwide rice production and hence food security has been threatened by a number of diseases of viral, bacterial and fungal origin. At present out of 70 known diseases of rice only 11 are of bacterial origin [3]. Among these diseases, bacterial blight (BB) is considered as one of the oldest and fourth major known diseases of rice [4,5]. High pathogenicity of bacterial blight pathogen gives it paramount importance globally and considered as a major hindrance towards acquiring sustainable rice productivity.

*Xanthomonas oryzae* pv. *oryzae*, a gram-negative bacterium, is responsible for BB disease. Though it was first reported in Fukuoka, Kyushu, Japan in 1884, it became widespread in Southeast Asia during 1960s after the introduction of high-yielding varieties [6,7]. As one of the major devastating rice diseases, bacterial blight can lead up to 50% rice yield loss [8], which leads to 16% of crop yield loss globally [9]. In Bangladesh yield loss due to bacterial blight ranges from 5.8 to 30.4% depending on growth stage, variety and environmental conditions [10].

Sustainable management of rice bacterial blight disease has been a major global concern. Deployment of cultivar with resistant gene (s) would be an environmentally safe, economic and effective approach to control bacterial blight disease and curtail yield loss [11,12]. So far, around 45 resistance (*R*) genes against BB have been identified in diverse rice sources [13].

Nevertheless, only 11 of these *R* genes (*Xa1*, *Xa3*/*Xa26*, *Xa4*, *xa5*, *xa13*, *Xa21*, *Xa23*, *Xa25*, *Xa27* and *Xa41*) have been cloned and functionally assessed. Based on the proteins encoded by the cloned-*R* genes and mode of action they are further classified into four groups: RLK (Receptor-like kinase) genes (*Xa4*, *Xa21* and *Xa3*/*Xa26*), SWEET (Sugar will eventually be exported transporter) genes (*xa13*, *xa25* and *xa41*), executor genes (*Xa10*, *Xa23* and *Xa27*) and other types of genes (*Xa1* and *xa5*) [11]. All these cloned R genes and some of the un-cloned *R* genes, like *Xa7*, are widely utilized in rice breeding programs especially focused on pyramiding of multiple resistant genes to develop bacterial blight resistant rice varieties. However, wild rice species and germplasm collections are widely considered as a rich reservoir of valuable genes and so, exploring germplasm collection in search of resistant cultivar has become a major focus of the researchers [14-17].

Nonetheless, epistasis and masking of gene effect rendered the conventional approach ineffective in identifying cultivars carrying multiple disease resistant genes [18]. Alternatively, molecular approach is widely accepted and utilized as a highly reliable and efficient method in identifying resistant genes [12,19,20]. Interestingly, in a recent pathotype profiling study conducted on 118 *Xoo* isolates of Bangladesh, we identified *xa5*, *xa8*, *xa13*, *Xa21* and *Xa23* genes as the most effective *R* genes against diverse *Xoo* strains [21]. In line with our previous finding, we aimed to identify five cloned *R* genes viz. *Xa4*, *xa5*, *xa13*, *Xa21*, *Xa23* and one un-cloned but widely utilized *R* gene, *Xa7*, from a local germplasm collection of Bangladesh using STS markers.

On the other hand, knowledge of genetic diversity of germplasm collection helps to attain durable selection gain while, plays a major role in parent identification and thus increases the likelihood of crop improvement [22-24]. Then again, molecular markers play a leading role in genetic diversity study. A wide range of PCR (Polymerase Chain Reaction) based molecular markers like, simple sequence repeat (SSR), inter-simple sequence repeat (ISSR), restriction fragment length polymorphism (RFLP), amplified fragment length polymorphism (AFLP), sequence-related amplified polymorphism (SRAP) are currently being deployed for genetic diversity assessment [25,26]. Among them dominant marker ISSRs, developed by [27], are simple, fast, cost effective, highly reproducible and are widely utilize molecular marker for genetic diversity assessment in plants [28-31]. Nevertheless, rice cultivars are often classified based on quantitative traits. So, by combining molecular and morphological approach will be a more comprehensive strategy to assess genetic divergence and to ensure a precise genotype documentation along with parental line selection for breeding programs.

In the current study, we explored 792 landraces of Bangladesh through molecular and morphological approaches with an objective to identify bacterial blight resistant germplasm and evaluate genetic diversity among them to enrich and diversify the rice gene pool for future breeding programs. The outcome of this study will be useful for research on rice and rice improvement programs.

## Materials and Methods

### Collection of germplasm and experimental design

A total of 792 germplasm (S1 Table) along with six bacterial blight resistant *R* gene containing monogenic lines viz., IRBB4 (*Xa4*), IRBB5 (*xa5*), IRBB7 (*Xa7*), IRBB13 (*xa13*), IRBB21 (*Xa21*), IRBB23 (*Xa23*), one susceptible line IR24, one pyramided resistant line IRBB60 (*Xa4*, *xa5*, *xa13* and *Xa21*) and one elite variety BRRI dhan28 were collected from Genetic resource and seed division (GRSD), Bangladesh Rice Research Institute (BRRI), Gazipur, Bangladesh and International Rice Research Institute (IRRI), Philippines. All 792 germplasm were Bangladeshi landraces. In order to evaluate bacterial blight resistance, seeds of all germplasm along with the BRRI dhan28 and IRBB60 were sown in a seedling nursery. Twenty-one days old seedling were transplanted to the experimental field of Plant Pathology Division, BRRI, Gazipur (Longitude: 23.9903° N, Latitude: 90.3996° E) following randomized complete block design (RCBD) with three replications.

### Inoculum preparation and pathogenicity test

Three major highly virulent strains of *Xanthomonas oryzae* pv. *oryzae* viz., *BXo69*, *BXo87* and *BXo93* [21] were collected from Plant Pathology Division, BRRI, Gazipur and used for pathogenicity test. Bacterial (*Xoo*) inoculums were grown on peptone sucrose agar (PSA) medium at 28℃ for 48 hours. The inoculum was diluted with distilled water and the absorbance was adjusted to approximately OD_600_ = 1. This value corresponds to the concentration of about 3.3×10^8^ colony forming units per milliliter (CFU/mL), which normally provides optimum *Xoo* infection in the host.

Rice plants were inoculated at maximum tillering stage by following leaf clipping method [32]. After inoculation plants were monitored in every 24-h interval to note disease appearance. The disease severity data (Lesion length) were recorded at 14 days of inoculation from 10 leaves of each entry. On the basis of disease severity, entries were classified as highly resistant <1 cm (Score 0), resistant 1.1-3 cm (Score 1), moderately resistant 3.1-5 cm (Score 3), moderately susceptible 5.1-10 cm (Score 5), susceptible 10.1-15 cm (Score 7) and highly susceptible >15 cm (Score 9) [33]. The lesion length covering the whole infected region of the leaf was measured with a scale. For our further study of resistant gene detection and genetic diversity analysis, we selected highly resistant to moderately resistant (Score 0-3) 74 germplasm based on the pathogenicity test.

### DNA extraction and PCR amplification

Fresh, young leaves were collected and stored in 1.5 mL microcentrifuge tubes at -20℃. DNA extraction was carried out following modified CTAB (hexadecyltrimethylammonium bromide) method [34]. Extracted DNA pellets were dissolved in 100μl of 1X TE (Tris EDTA) buffer. DNA quality and concentration were checked using NanoDrop spectrophotometer (Jenova Nano, UK). Finally, the working DNA solution was prepared by diluting stock solution to 25-30 ng/μl DNA concentration using 1X TE buffer and stored at 4℃.

Polymerase chain reaction (PCR) were performed to identify resistance gene(s) and determine genetic diversity among the selected germplasm. For the detection of bacterial blight resistant gene(s) six co-segregating STS markers tightly linked to *Xa4*, *xa5*, *Xa7*, *xa13*, *Xa21*, *Xa23* genes were used (Table 1). On the contrary, genetic diversity of the selected germplasm were determined using ISSR marker. A total of 22 ISSR markers were taken for polymorphism study and finally 17 markers were selected based on their unambiguous reproducible polymorphic band generation capability for genetic diversity analysis. PCR reaction mixture was formulated using 5 μl GoTaq G2 green PCR master-mix (Promega, USA), 1 μl primer (forward and revers, 0.5 μl each), 2 μl ddH_2_O and 2 μl genomic DNA solution. The PCR program involved: 4-minute initial denaturation step at 94°C, 35 cycles of denaturation at 94℃ for 30 second, annealing at 45℃ to 58℃ for 30 second and elongation at 72℃ for 1 minute to 1 minute 40 second, followed by a final elongation phase at 72℃ for 7 minutes.

**Table 1:**
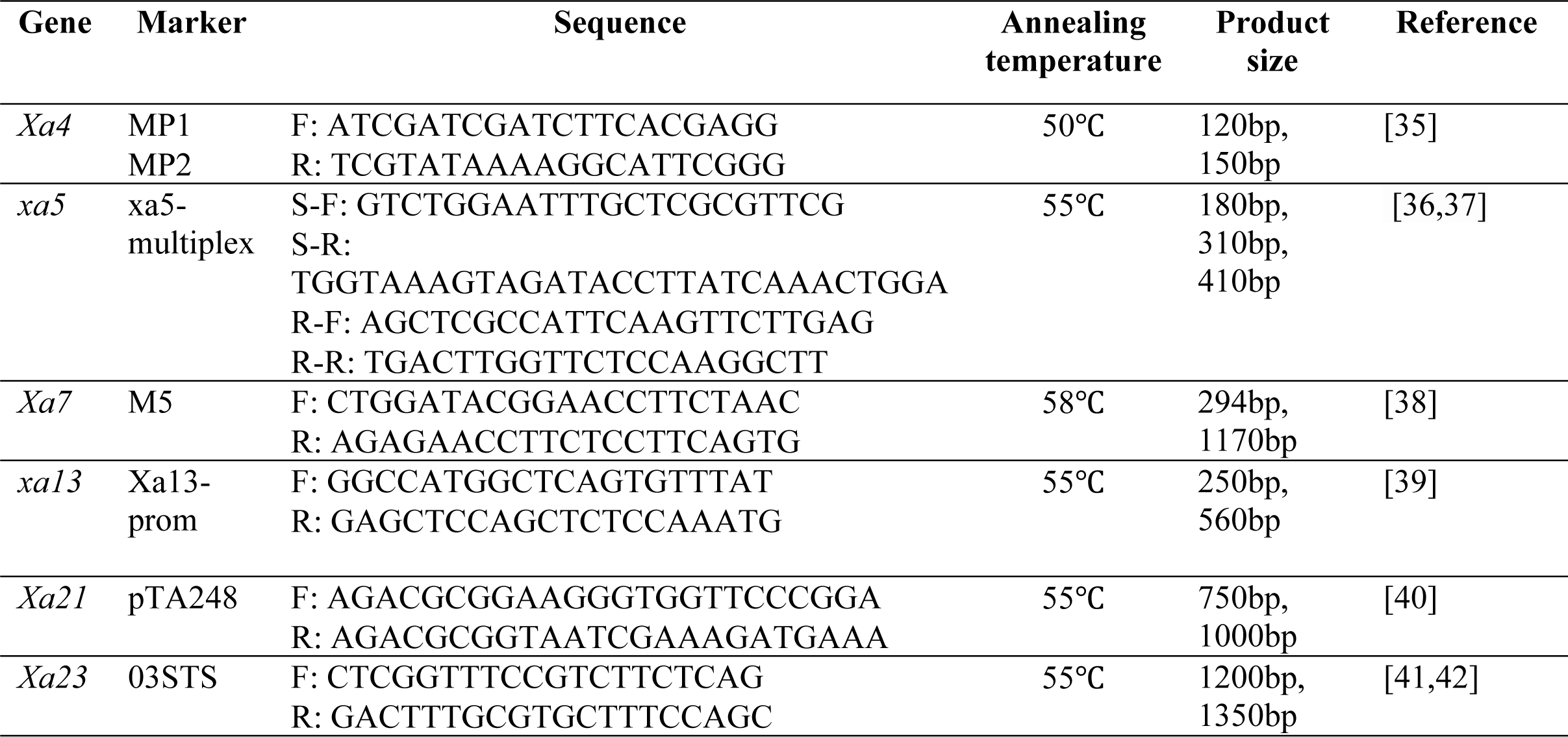
List of primers used for resistant detection with their sequence, annealing temperature and expected product size.

The PCR product of *Xa4* and *xa5* was resolved in 2.5% agarose gel while, the PCR product of *Xa7*, *xa13*, *Xa21*, *Xa23* were resolved in 1.5% agarose gel using 1X TBE buffer at 90 V-120 minutes. Monogenic resistant line of the respective gene was used as resistant check and IR24 was used as susceptible check for identification of the resistant genes. On the other hand, PCR product of all ISSR markers were resolved in 10% (w/v) polyacrylamide gel in 1X TBE at 90 V-70 min. For visualization of the resolved DNA band all gels were stained with ethidium bromide (10 μg/mL) solution and finally visualized under transilluminator (UVP GelSolo).

### Characterization of the quantitative traits

Quantitative traits of the74 bacterial blight resistant germplasm were also evaluated. Twenty-five days old seedlings were transplanted following RCB design with three replications in the experimental field of Plant Pathology Division, BRRI, Gazipur with a spacing of 20cm × 20cm. Quantitative trait data such as, days to 50% flowering, days to maturity, plant height (cm), number of tiller hill^-1^, number of effective tiller hill^-1^, panicle length (cm), spikelet panicle^-1^, filled spikelet panicle^-1^, unfilled spikelet panicle^-1^ and 1000 grain weight (g) were documented with three replications.

### Data analysis

ISSR band profile of each marker was scored using binary system where, score 1 indicated presence and score 0 indicated absence of a band. A UPGMA dendrogram was constructed using Jaccard’s similarity coefficient using NTSYS-pc version 2.21q software and two dimensional PCA (Principe component analysis) scattered plot was constructed using R-4.1.0 software. Polymorphic information content (PIC) value of the markers was calculated following the formula of [43] while, resolving power (RP) of each ISSR marker was calculated using the method proposed by [44].

For quantitative trait data, analysis of variance (ANOVA) and mean comparison test (LSD) were performed using Statistix 10 software. NTSYS-pc version 2.21q software was used to construct Euclidean distance based UPGMA dendrogram while two dimensional PCA scattered plot, correlogram based on Pearson’s correlation among the quantitative traits and a heatmap based on trait mean values of different clusters of UPGMA dendrogram were constructed using R-4.1.0 software. Mantel test was performed using NTSYS-pc version 2.21q software to find correlation between molecular and morphological data matrix.

## Results

### Identification of bacterial blight resistant germplasm

Based on disease reaction pattern of 792 germplasm against the three virulent strains of *Xoo*, only one germplasm was found highly resistant (score 0), 33 germplasm were resistant (score 1) and 40 germplasm were found moderately resistant (score 3) against three virulent strains (Fig 1). Rest of the germplasm showed moderately susceptible to highly susceptible disease reaction (S1 Table). IRBB60 showed highly resistant score (0) while BRRI dhan28 showed highly susceptible reaction (score 9).

**Fig 1.**
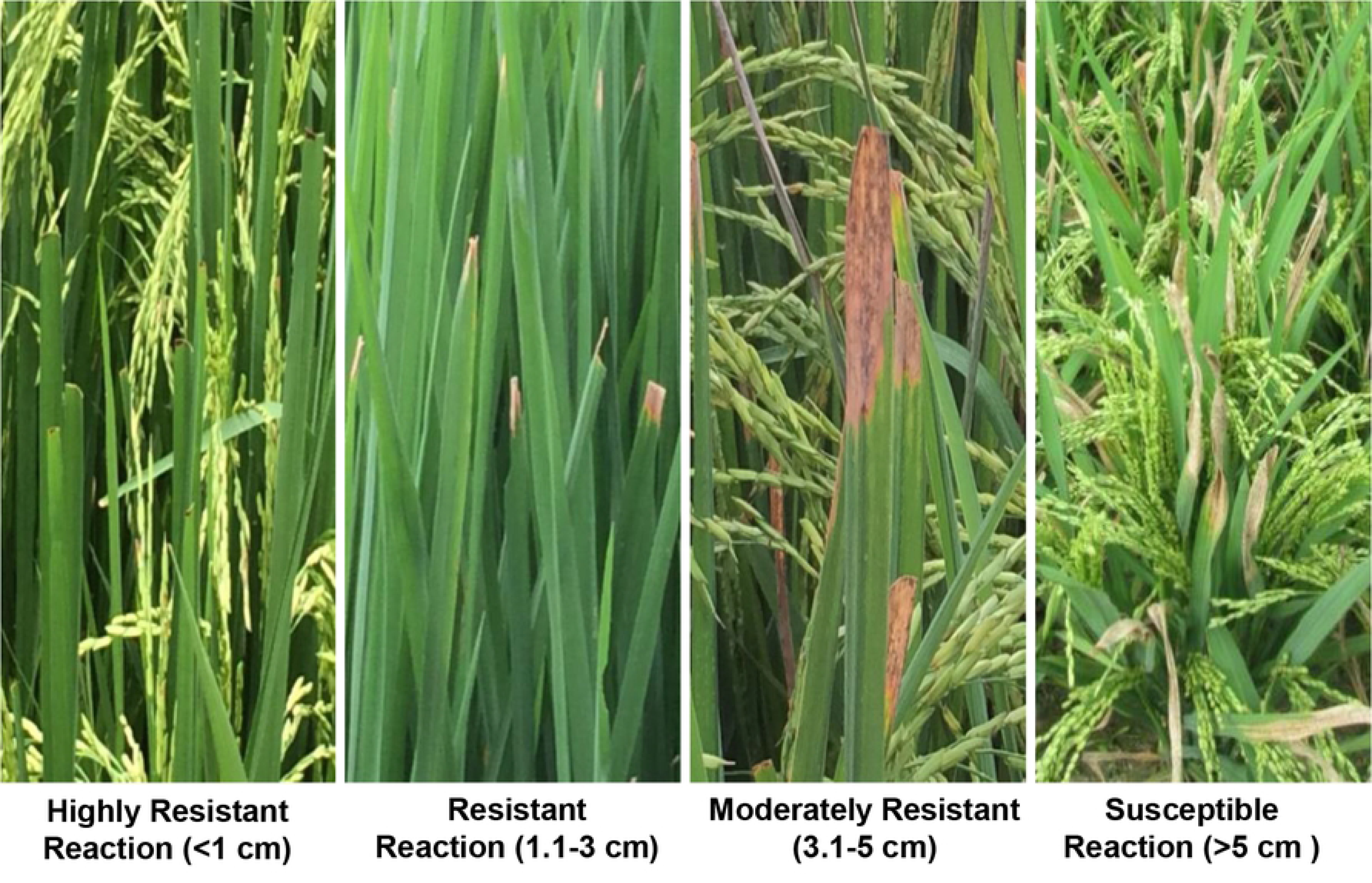
Plants with different lesion length caused by *Xanthomonas oryzae* pv*. oryzae* and their resistance status

### Identification of the resistant genes

MP1-2 (*Xa4* gene linked) primer detected 41 germplasm (55.41%) as *Xa4* positive and 33 germplasm (44.59%) as *Xa4* negative (Fig 2A). On the other hand, xa5-multiplex primer (*xa5* linked) identified 15 germplasm (20.27%) harboring *xa5* allele while, 59 germplasm (79.73%) were found negative (Fig 2B). Nonetheless, M5 primer (*Xa7* gene linked) distinguished 62 germplasm (83.78%) as *Xa7* positive while, the rest 12 (16.22%) were found negative (Fig 2C). Nevertheless, xa13-prom primer (*xa13* gene linked) detected 33 germplasm (44.59%) to carry *xa13* (recessive) allele while, 41 germplasm (55.41%) to carry *Xa13* (dominant) allele (Fig 2D). Notably, pTA248 primer (*Xa21* gene linked) dug out only a single germplasm (G3, Acc no.4216) carrying *Xa21* gene (Fig 2E) and finally, 03STS primer (*Xa23* gene linked) detected 19 germplasm (25.68%) as *Xa23* positive and 55 germplasm (74.32%) as *Xa23* negative (Fig 2F). A summary of the disease reaction score with gene identification of the 74 potential germplasm are shown in Fig 3 and S2 Table. Interestingly, most of the resistant germplasm were found to carry multiple genes of different combinations. Among the germplasm, 10 germplasm consisted of 4 genes, 15 germplasm consisted of 3 genes and 38 germplasm consisted of 2 genes of various combinations (Fig 4). Particularly, germplasm G3 (Acc. No. 4216; having *Xa4*, *Xa7*, *xa13* and *Xa21*) was only highly resistant germplasm among the 792 germplasm in the pathogenicity test.

**Fig 2.**
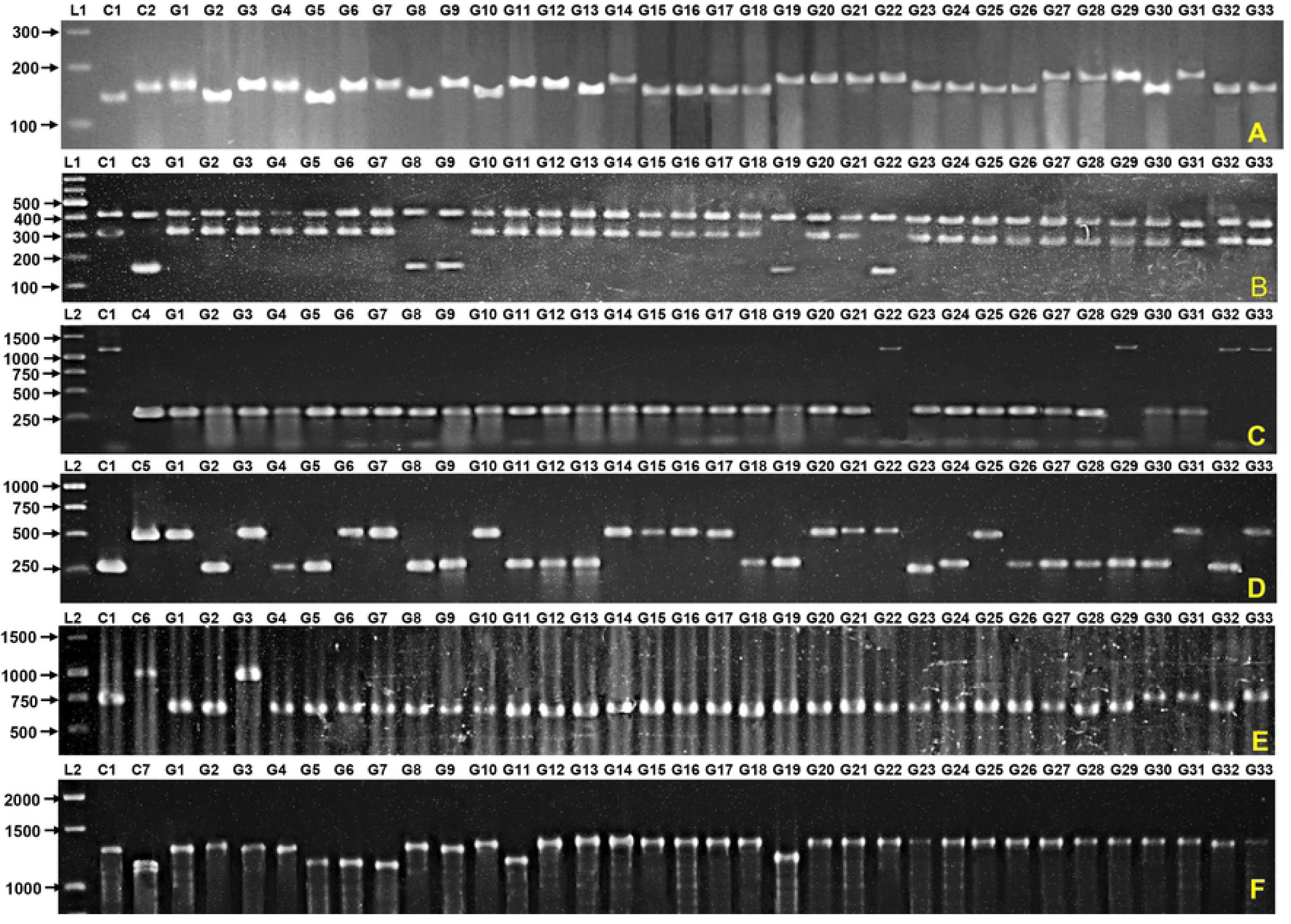
Partial gel picture of, (A) M1-2 primer (*Xa4* gene linked), (B) *xa5* multiplex primer (*xa5* gene linked), (C) M5 primer (*Xa7* gene linked), (D) xa13-prom primer (*xa13* gene linked), (E) pTA248 primer (*Xa21* gene linked), (F) 03STS primer (*Xa23* gene linked), resolved in 1.5-2% agarose gel. Here, C1- susceptible check IR24 and C2- IRBB4, C3- IRBB5, C4- IRBB7, C5- IRBB13, C6- IRBB21 and C7- IRBB23. L1 corresponds to 100bp DNA ladder and L2 corresponds to 1kb DNA ladder. G1 to G33 represents first 33 germplasm out of 74 potential germplasm identified based on the disease reaction against three *Xoo* strains.

**Fig 3.**
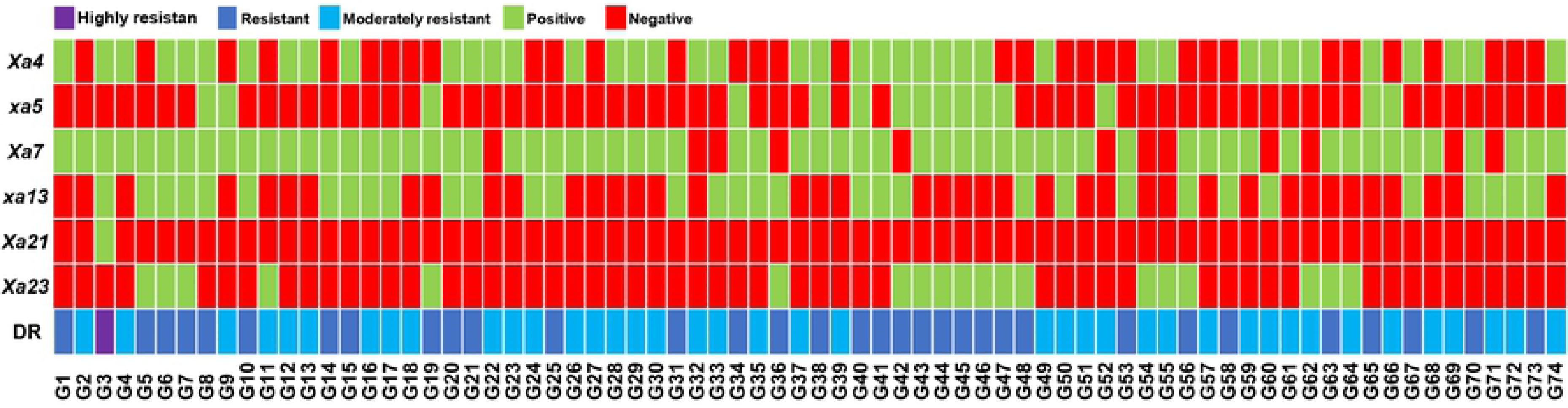
Color map representing disease reaction (DR) status of 74 germplasm against three *Xoo* strains and the resistant gene(s) contained by them. Green color indicated presence of a gene while red color indicated absence of a gene. Presence or absence of a gene is confirmed by STS marker analysis of the respected gene.

**Fig 4.**
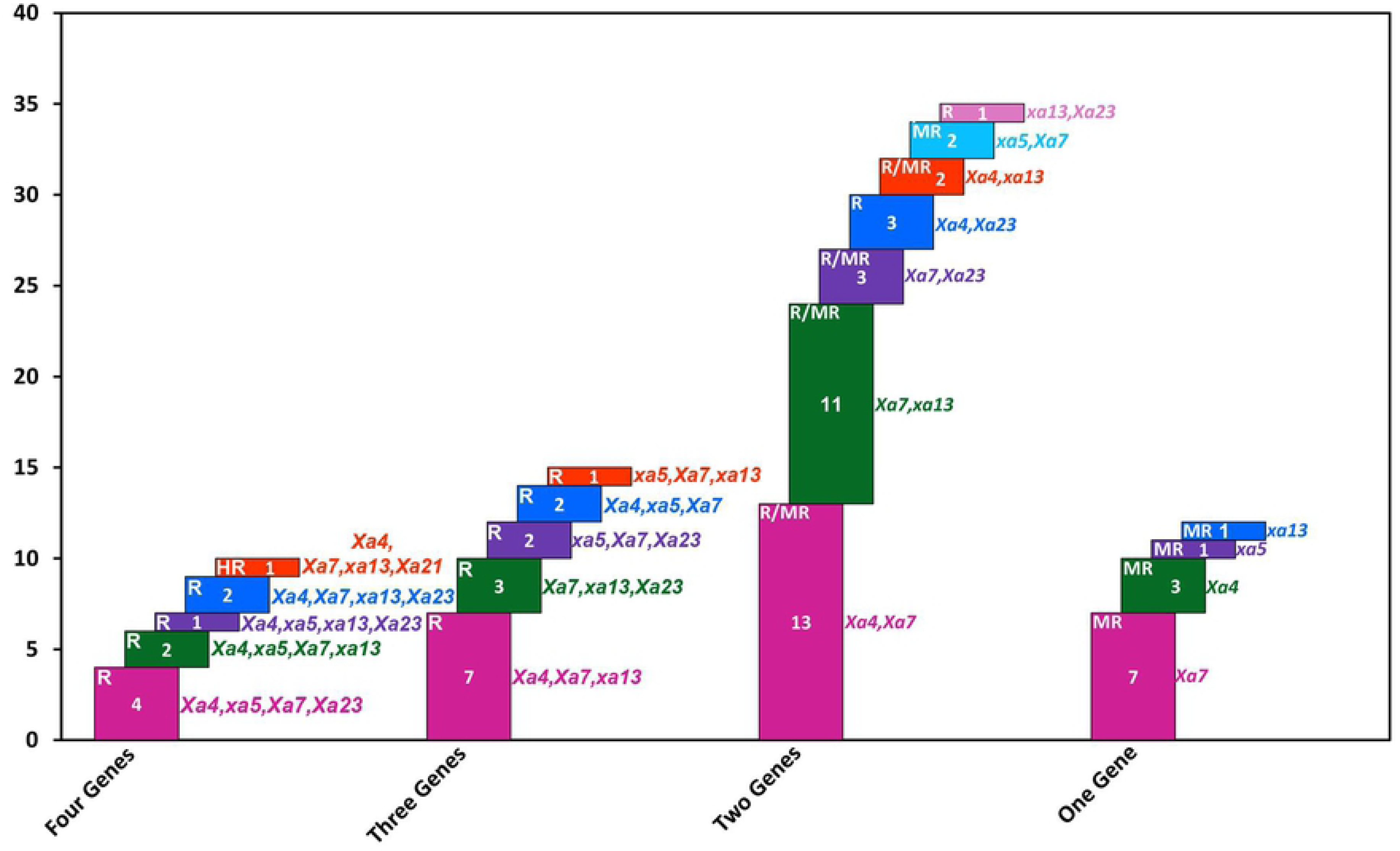
Different gene combinations recorded in the 74 selected germplasm and their resistant status in pathogenicity test. Each colored cell in a bar represents a different gene combination. Number inside the cell represents the frequency of that gene combination among 74 germplasm. Here, HR- highly resistant, R-resistant, MR-moderately resistant.

Notable, rest of the 4 to 3 genes carrying germplasm were found resistant and germplasm carrying 2 genes were found resistant to moderately resistant in the pathogenicity test. Germplasm with a single *R* gene were found moderately resistant. Markedly, germplasm G43 (Acc. No. 1523) was found to carry three of the *R* genes (*xa5*, *xa13* and *Xa23*) effective against diverse virulent *Xoo* strains of Bangladesh, along with *Xa4* gene.

### Genetic diversity of resistant germplasm based on molecular markers

Seventeen ISSR markers generated a total of 140 definite bands, out of which 105 bands (75%) were found polymorphic. The average number of polymorphic bands per loci were 6.8. The PIC value ranged from 0.16 to 0.49 with an average of 0.31 while the resolving power of the markers ranged from 0.20 to 0.89 with an average of 0.44 (Table 2). Partial DNA profile of some ISSR markers presented in Fig 5.

**Fig 5.**
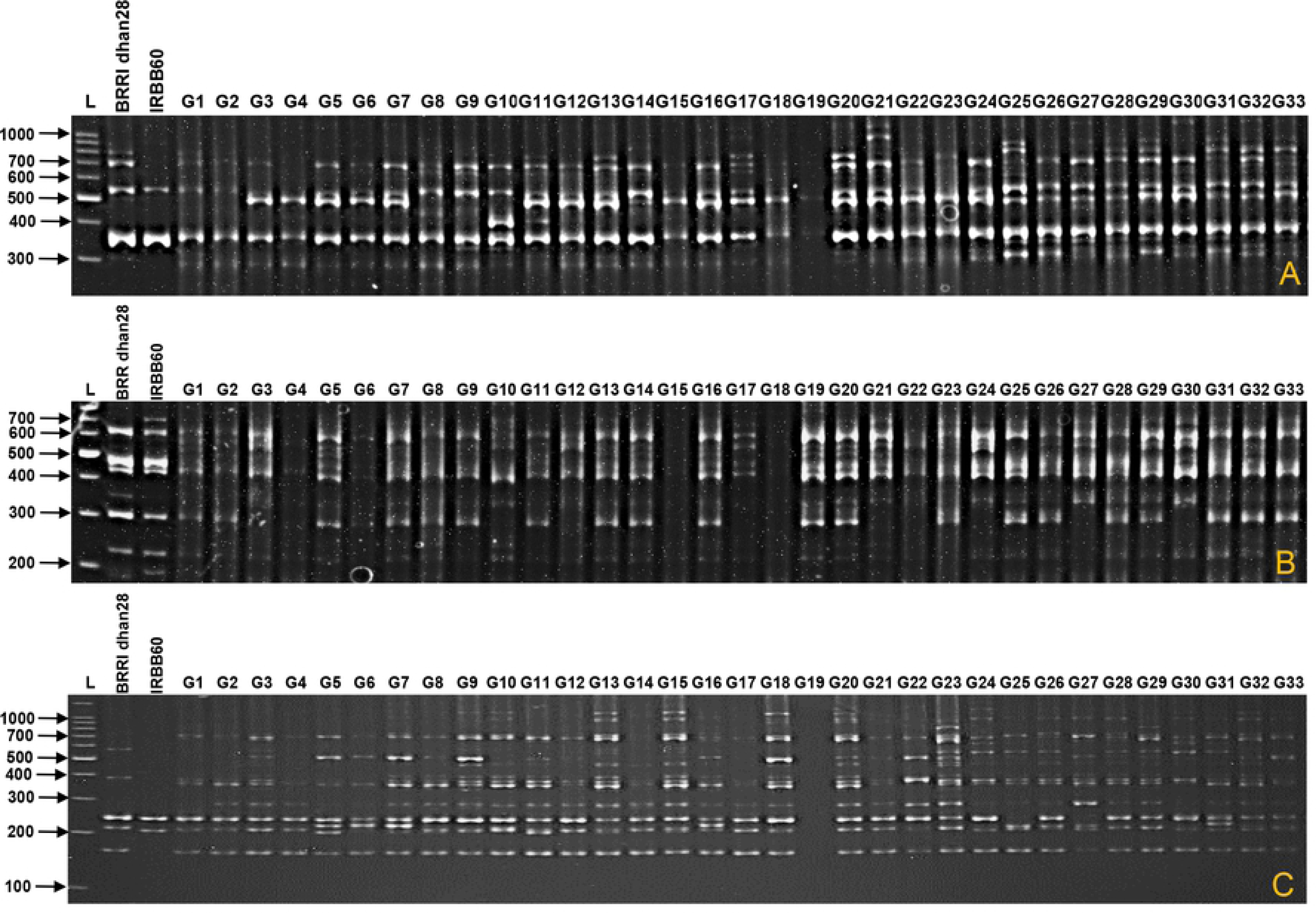
Partial gel picture of three ISSR markers, (a) ISSR808, (b) ISSR809, (c) ISSR810, resolved in 10% polyacrylamide gel. G1 to G33 represents first 33 germplasm out of 74 potential germplasm identified based on the disease reaction against three *Xoo* strains. L corresponds to 100bp DNA ladder.

**Table 2:** List of ISSR markers with their sequence, annealing temperature, number of polymorphic bands, PIC value and RP value.

| Marker<sup>(a)</sup> | Sequence | Annealing temperature | Polymorphic band | PIC | RP |
| --- | --- | --- | --- | --- | --- |
| UBC807 | (AG)<sub>8</sub> T | 45°C | 4 | 0.26 | 0.34 |
| UBC808 | (AG)<sub>8</sub> C | 45°C | 8 | 0.21 | 0.26 |
| UBC809 | (AG)<sub>8</sub> G | 45°C | 7 | 0.31 | 0.46 |
| UBC810 | (GA)<sub>8</sub> T | 45°C | 11 | 0.22 | 0.28 |
| UBC836 | (AG)<sub>8</sub> YA | 45°C | 6 | 0.16 | 0.24 |
| UBC840 | (GA)<sub>8</sub> YT | 45°C | 2 | 0.49 | 0.89 |
| UBC848 | (CA)<sub>8</sub>RG | 50°C | 2 | 0.30 | 0.41 |
| UBC853 | (TC)<sub>8</sub>RT | 45°C | 9 | 0.27 | 0.39 |
| UBC855 | (AC)<sub>8</sub>YT | 50°C | 5 | 0.30 | 0.41 |
| UBC857 | (AC)<sub>8</sub>YG | 50°C | 3 | 0.46 | 0.40 |
| UBC884 | HBH(AG)<sub>7</sub> | 45°C | 4 | 0.38 | 0.53 |
| UBC885 | BHB(GA)<sub>7</sub> | 45°C | 6 | 0.41 | 0.66 |
| UBC886 | VDV(CT)<sub>7</sub> | 45°C | 4 | 0.32 | 0.46 |
| UBC888 | BDB(CA)<sub>7</sub> | 50°C | 4 | 0.36 | 0.55 |
| UBC889 | DBD(AC)<sub>7</sub> | 50°C | 13 | 0.34 | 0.51 |
| UBC890 | VHV(GT)<sub>7</sub> | 50°C | 11 | 0.36 | 0.53 |
| UBC891 | HVH(TG)<sub>7</sub> | 50°C | 6 | 0.18 | 0.20 |
| Total | — | — | 105 |  |  |
| Mean | — | — | 6.18 | 0.31 | 0.44 |

Note: (a) UBC primer set 9, University of British Columbia. PIC: Polymorphic Information Content, RP: Resolving Power.

UPGMA dendrogram, based on the binary data of ISSR markers grouped the 74 landraces, IRBB60 and BRRI dhan28 into 20 groups at the coefficient value of 0.135 where, Jaccard’s coefficient value ranged from 0 to 0.525 (Fig 6). Out of 20 clusters, cluster-V (25 germplasm), cluster-X (11 germplasm), cluster-IV (10 germplasm), cluster-XVI (6 germplasm), cluster-IX (4 germplasm), cluster-XV (3 germplasm) and cluster-XVII (3 germplasm) were the larger clusters while, rest of the 12 clusters comprised of only one to two germplasm. Conversely, two-dimensional scattered plot generated from PCA analysis generated 16 major clusters (Fig 7) where, cluster-I contained the highest 28 germplasm followed by cluster-XI (11 germplasm), cluster-XIV (11 germplasm), cluster-XV (5 germplasm), cluster-XVI (5 germplasm), cluster-XII (4 germplasm) and cluster-IV (3 germplasm). The remaining clusters contained only one germplasm. Notably, both UPGMA dendrogram, PCA analysis pointed that some germplasm such as, G6, G15, G18, G25, G34, G49, G64 and G76 preferred to remain in a single group. However, according to the PCA analysis, 71.14% variability was explained by first five components where they individually accounted for 31.84%, 16.76%, 9.21%, 7.88% and 5.45 % variability.

**Fig 6.**
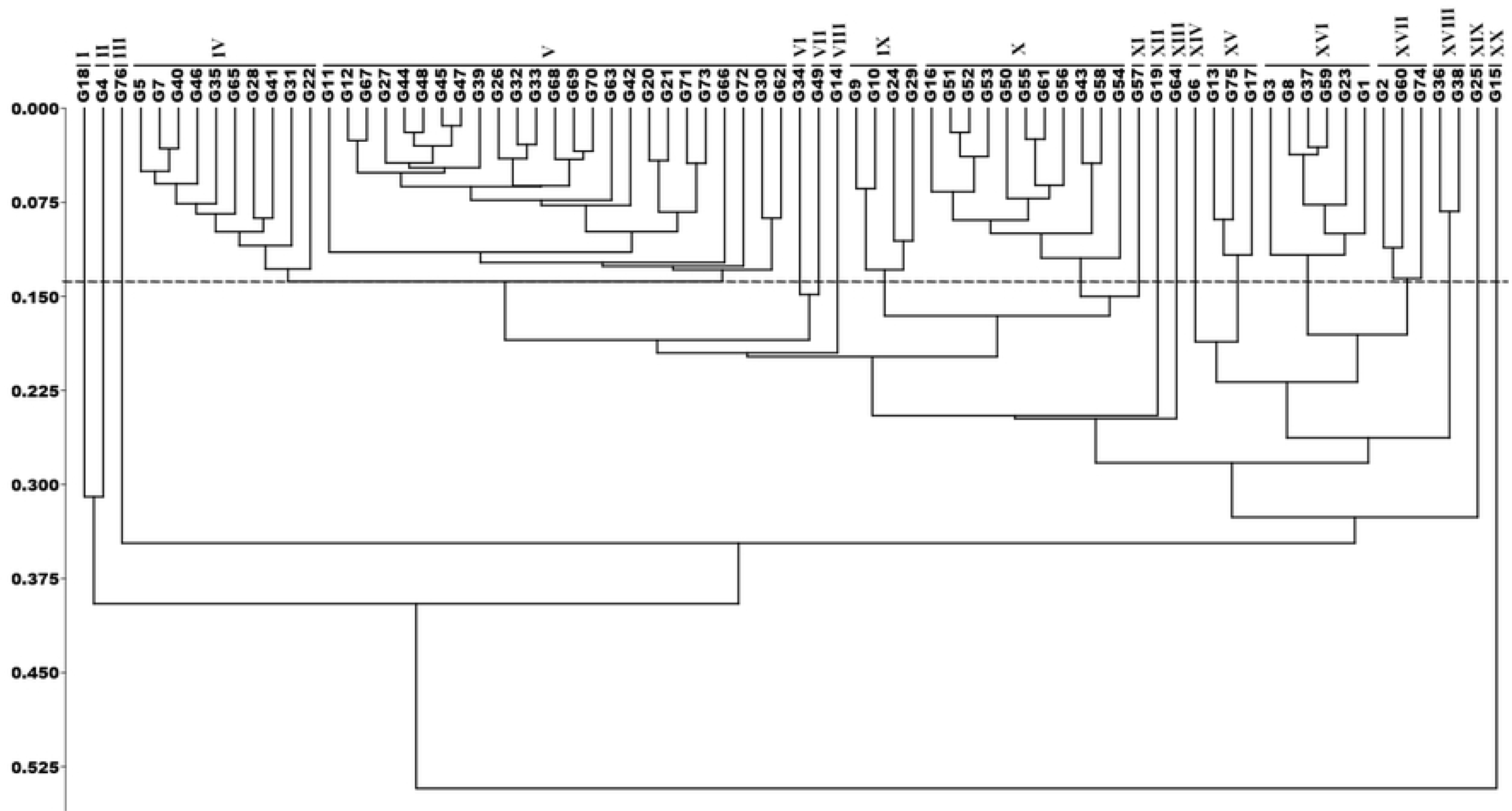
UPGMA dendrogram of 76 genotypes constructed from ISSR marker data based on Jaccard’s coefficient. Here, G1 to G74 represents 74 potential germplasm identified based on the disease reaction against 3 *Xoo* strains while G75 represents BRRI dhan28 and G76 represents IRBB60.

**Fig 7.**
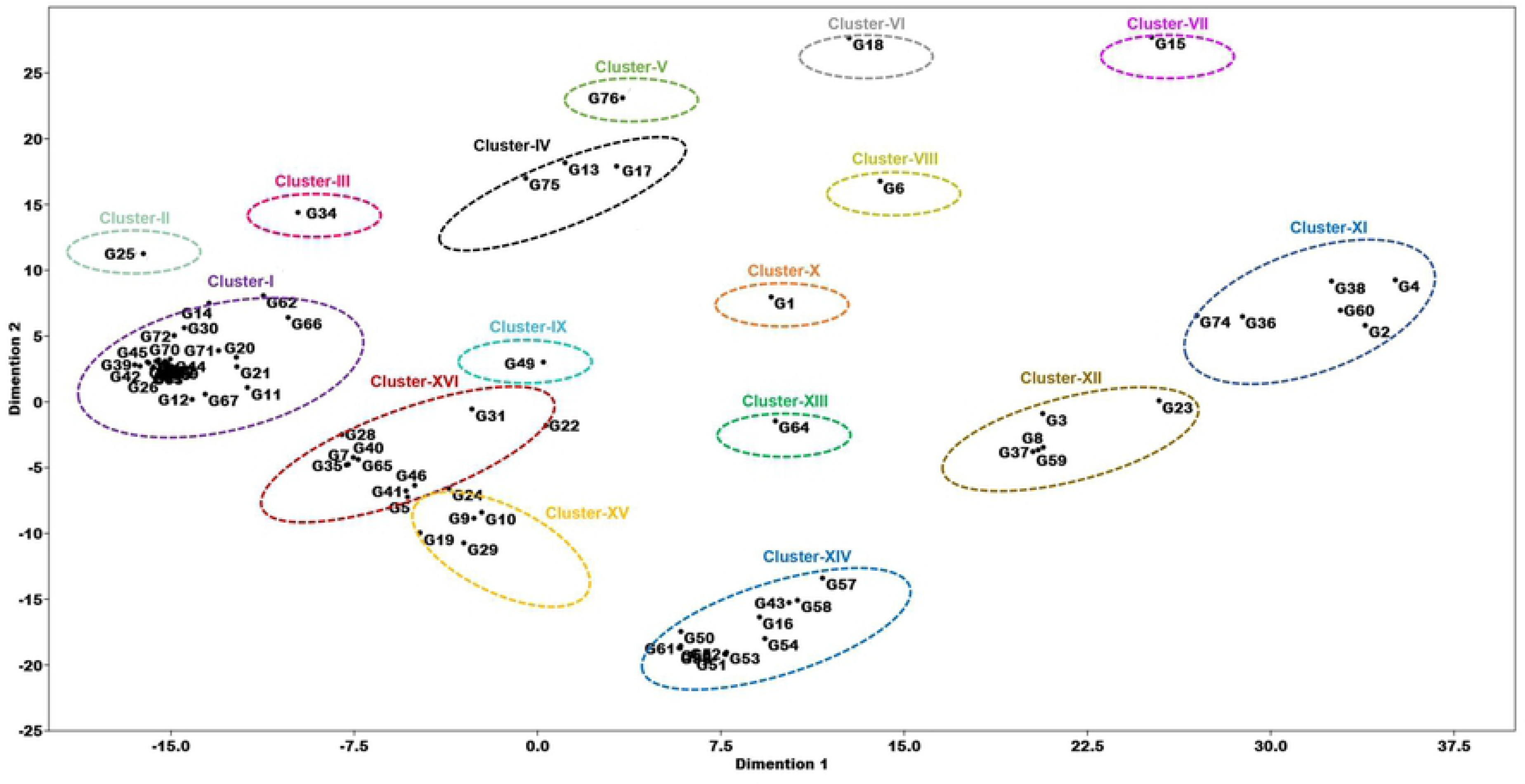
Two dimensional PCA scattered plot constructed from ISSR marker data grouping 76 genotypes into different clusters. Each different colored ellipse represents a different cluster. Here, G1 to G74 represents 74 potential germplasm identified based on the disease reaction against 3 *Xoo* strains while G75 represents BRRI dhan28 and G76 represents IRBB60.

Based on Jaccard’s similarity matrix highest genetic similarity existed between the germplasm G45-G46 (0.92) followed by, G65- G66 (0.91) and G9- G8 (0.90) (S3 Table). Contrariwise, lowest genetic similarity was found between G15- G76 (0.1) followed by G18- G76 (0.15), G4- G76 (0.25), G72- G15 (0.26), G38- G76 (0.27), G74- G76 (0.27), G12- G76 (0.28), G64- G76 (0.28), G4- G54 (0.28), G63- G76 (0.29), G4- G53 (0.29), G53- G54 (0.29). On the other hand, our bacterial blight susceptible elite variety BRRI dhan28 showed highest genetic similarity with germplasm G61 (Acc. No. 3101; similarity value 71) and lowest genetic similarity with germplasm G20 (Acc. No. 4004; similarity value 0.3) followed by G17 (Acc. No. 3981; similarity value 0.37) and G6 (Acc. No. 991; similarity value 0.38).

### Morphological variability among the germplasm

Analysis of variance (Table 3) revealed that all the selected germplasm had highly significant variability (*p*≤0.01) in terms of 10 quantitative traits (S4 Table). Pearson’s correlation among the quantitative traits indicated that there was a significant positive correlation between number of tiller hill^-1^ and number of effective tiller hill^-1^ (0.96), days to maturity and days to flowering (1.0), filled spikelet panicle^-1^ and spikelet panicle^-1^ (0.53), unfilled spikelet panicle^-1^ and spikelet panicle^-1^ (0.69) (Fig 8). On the contrary, a significant negative correlation was observed between panicle length and number of tiller hill^-1^ (-0.29), panicle length and number of effective tiller hill^-1^, unfilled spikelet panicle^-1^ and filled spikelet panicle^-1^ (-0.24), 1000 grains weight and unfilled spikelet panicle^-1^ (-0.24).

**Fig 8.**
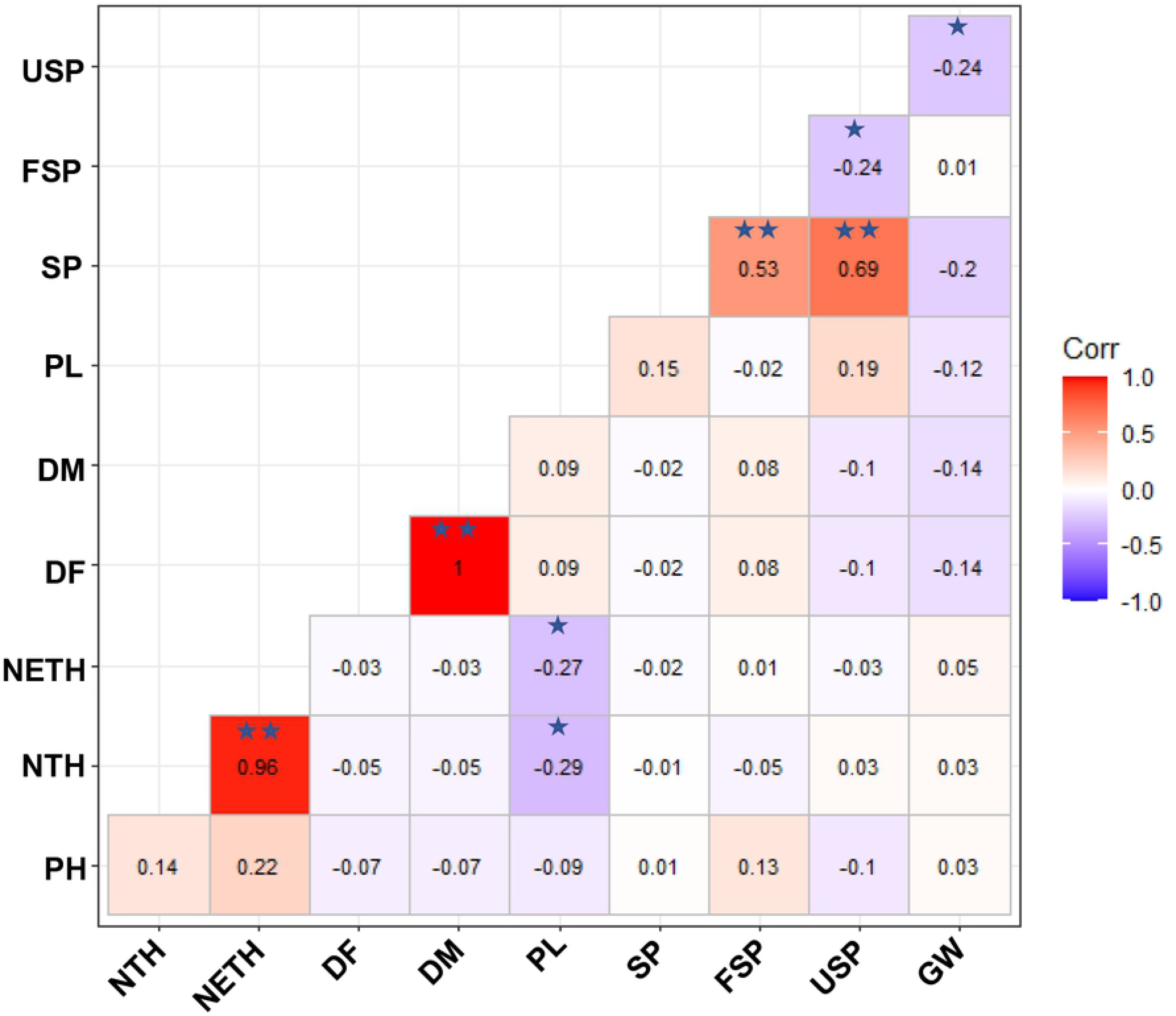
Correlogram presenting Pearson’s correlation between the quantitative traits. Red color represents positive correlation and violet color represents negative correlation. Here, PH-Plant height, NTH- Number of tiller hill^-1^, NETH- Number of effective tiller hill^-1^, DF- Days to flowering, DM- Days to maturity, PL- Panicle length, SP-Spikelet panicle^-1^, FSP- Filled spikelet panicle^-1^, USP- Unfilled spikelet panicle^-1^, GW- 1000 Grain weight. * p≤0.05, **p≤0.01.

**Table 3:**
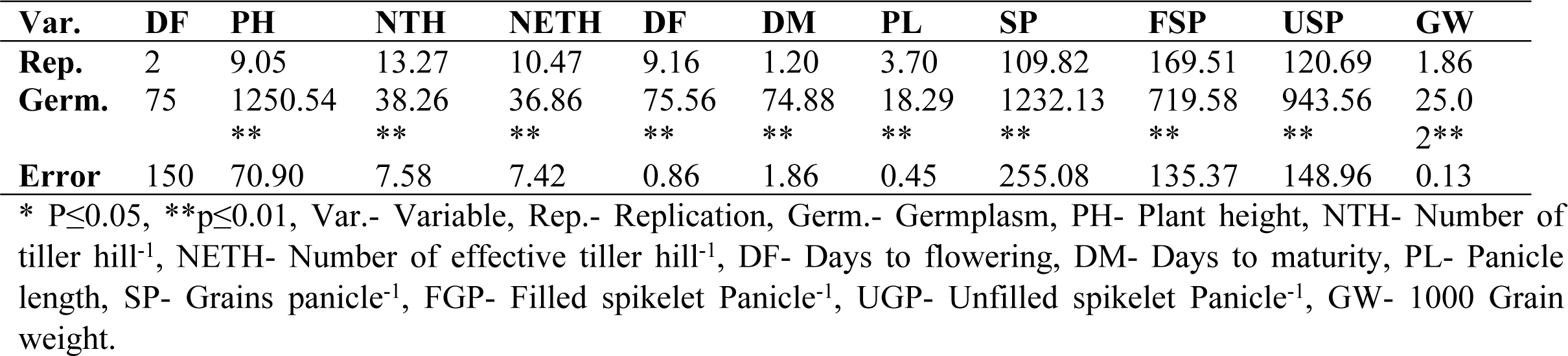
Mean squares values from analysis of variance for 10 quantitative traits among 74 rice germplasm with two check varieties.

Next, the UPGMA dendrogram based on Euclidean distance grouped the 74 germplasm into 8 different clusters where, cluster-V contained highest 18 germplasm followed by cluster-II (15 germplasm), cluster-VIII (9 germplasm), cluster-IV (8 germplasm), cluster-I, -III and -VII (7 germplasm) and finally cluster-VI (5 germplasm) (Fig 9). This clustering pattern was further supported by two-dimensional scattered plot from PCA analysis (Fig 10). In the PCA analysis 10 clusters were found, where 13 germplasm grouped into cluster-III and cluster-IV followed by cluster-I (10 germplasm), cluster-V and cluster-IX (9 germplasm), cluster-VIII (7 germplasm), cluster-II and cluster-VII (5 germplasm), cluster-X (3 germplasm) and cluster-VI (2 germplasm). According to the PCA analysis, 94.049% variability was observed due to first three component where they individually accounted for 43.294%, 30.13% and 20.625% variability.

**Fig 9.**
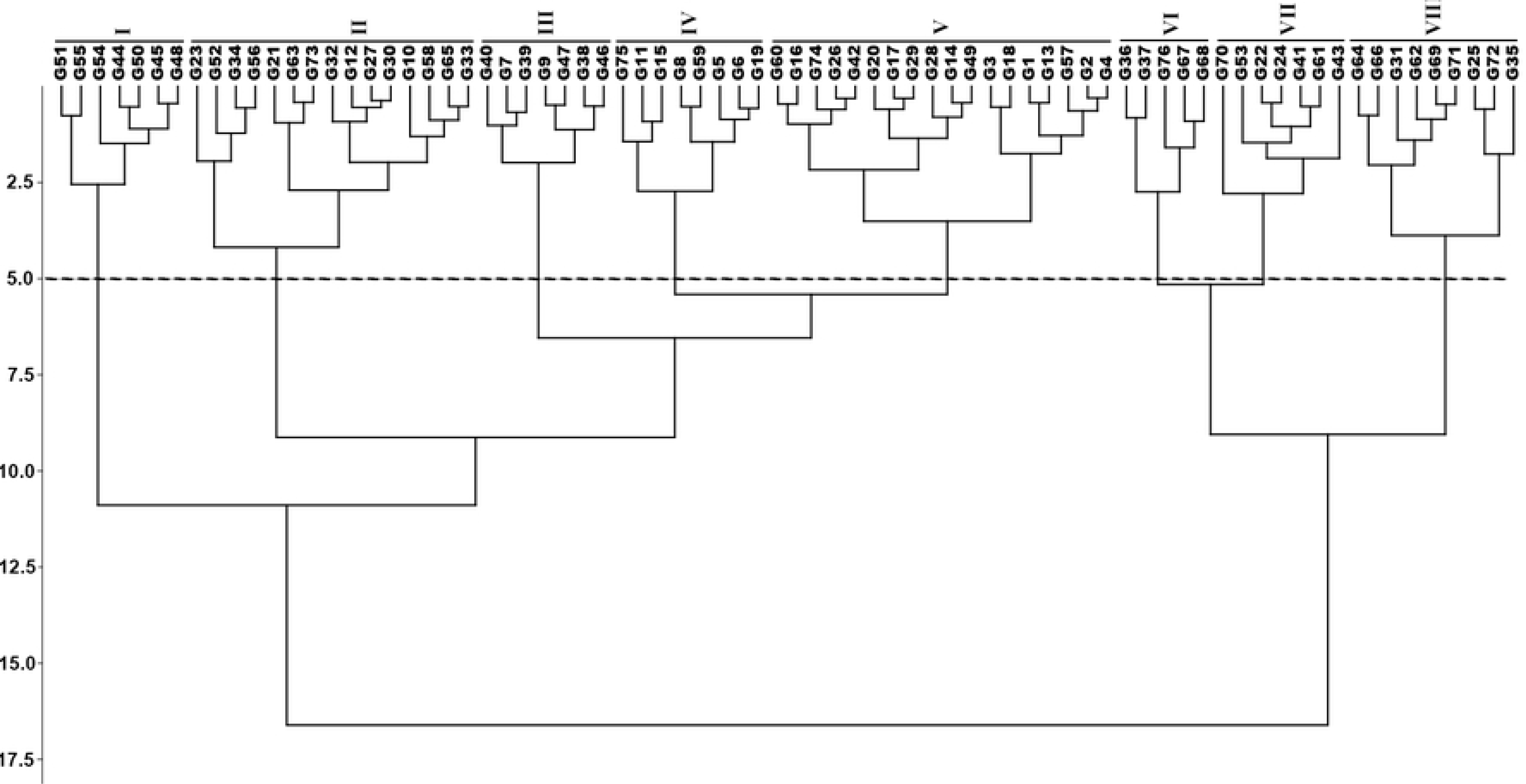
UPGMA dendrogram of 76 genotypes constructed from quantitative traits data based on Euclidean distance matrix. Here, G1 to G74 represents 74 potential germplasm identified based on the disease reaction against three *Xoo* strains while G75 represents BRRI dhan28 and G76 represents IRBB60.

**Fig 10.**
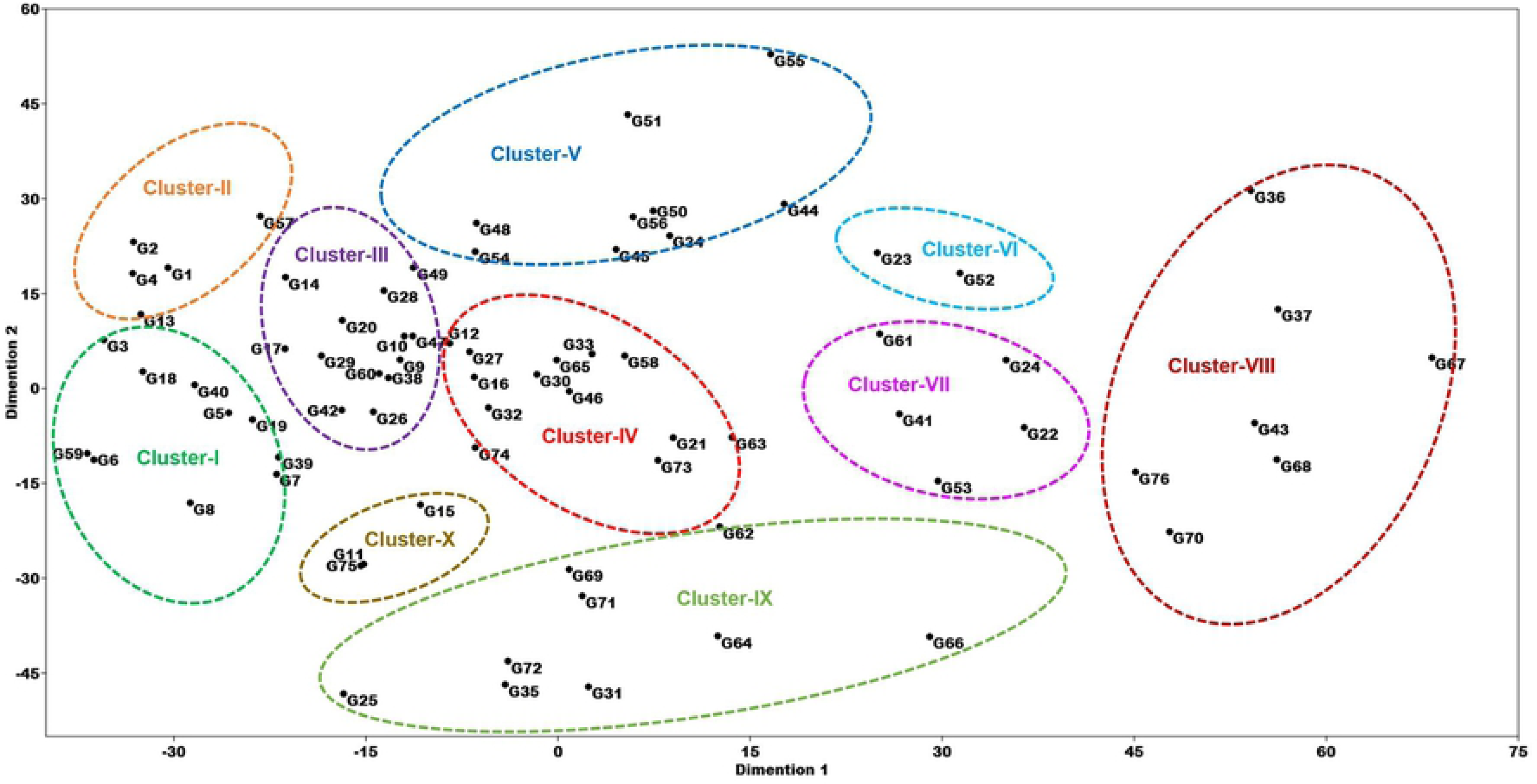
Two dimensional PCA scattered plot constructed from quantitative traits data grouping 76 genotypes into different clusters. Each different colored ellipse represents a different cluster. Here, G1 to G74 represents 74 potential germplasm identified based on the disease reaction against three *Xoo* strains while G75 represents BRRI dhan28 and G76 represents IRBB60.

The clustered germplasm from UPGMA dendrogram showed a variable mean performance in respect to quantitative traits. However, none of the clusters was found to have all the desired traits in respect to mean values (Fig 11). Notably, germplasm from Cluster-I had the highest mean value in respect to plant height, number of effective tiller hill^-1^ and filled spikelet panicle^-1^, while germplasm from cluster-II were recorded to have the lowest mean value in case of total tiller hill^-1^, effective tillers hill^-1^, days to flowering, days to maturity and unfilled spikelet panicle^-1^. However, the highest panicle length, unfilled spirelet panicle^-1^ along with the lowest 1000 grains weight in respect to mean performance were recorded in the germplasm from cluster-V. Nonetheless, the lowest mean plant height with the highest number of total tiller hill^-1^ were recorded in the germplasm from cluster-VIII.

**Fig 11.**
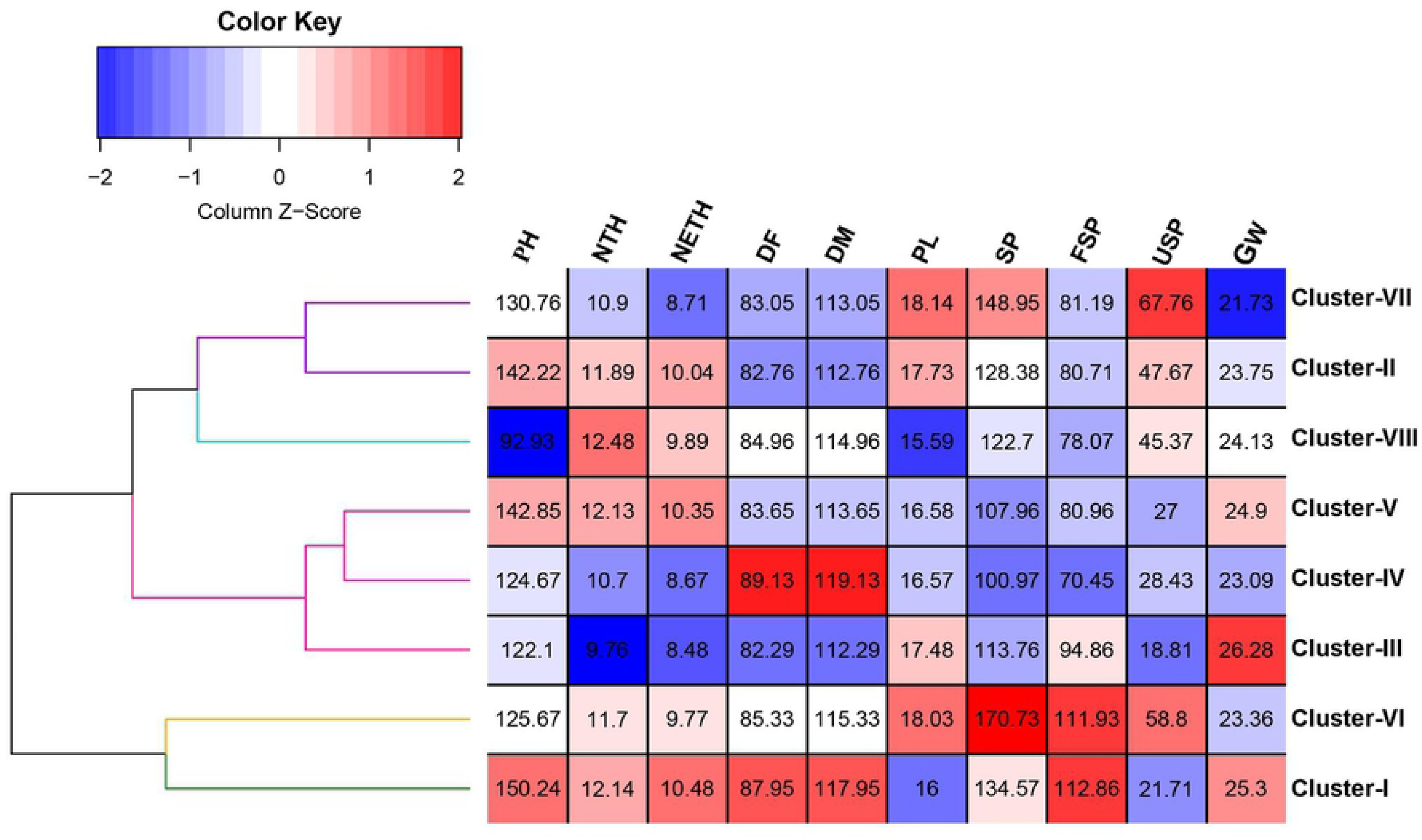
Heatmap showing quantitative trait performance in respect to mean value of different clustered germplasm grouped based on the UPGMA dendrogram constructed using morphological trait data. Blue color represents lower value while red color represents higher value. Here, PH- Plant height, NTH- Number of tiller hill^-1^, NETH- Number of effective tiller hill^-1^, DF- Days to flowering, DM- to maturity, PL- Panicle length, SP- Spikelet panicle^-1^, FGP- Filled spikelet panicle^-1^, UGP- Unfilled spikelet Panicle^-1^, GW- 1000 Grain weight.

Mantel test was performed to find correlation between molecular and morphological data matrix (Fig 12). Our findings suggested that a positive but non-significant correlation (Matrix correlation r= 0.113) existed between molecular and morphological variability data.

**Fig 12.**
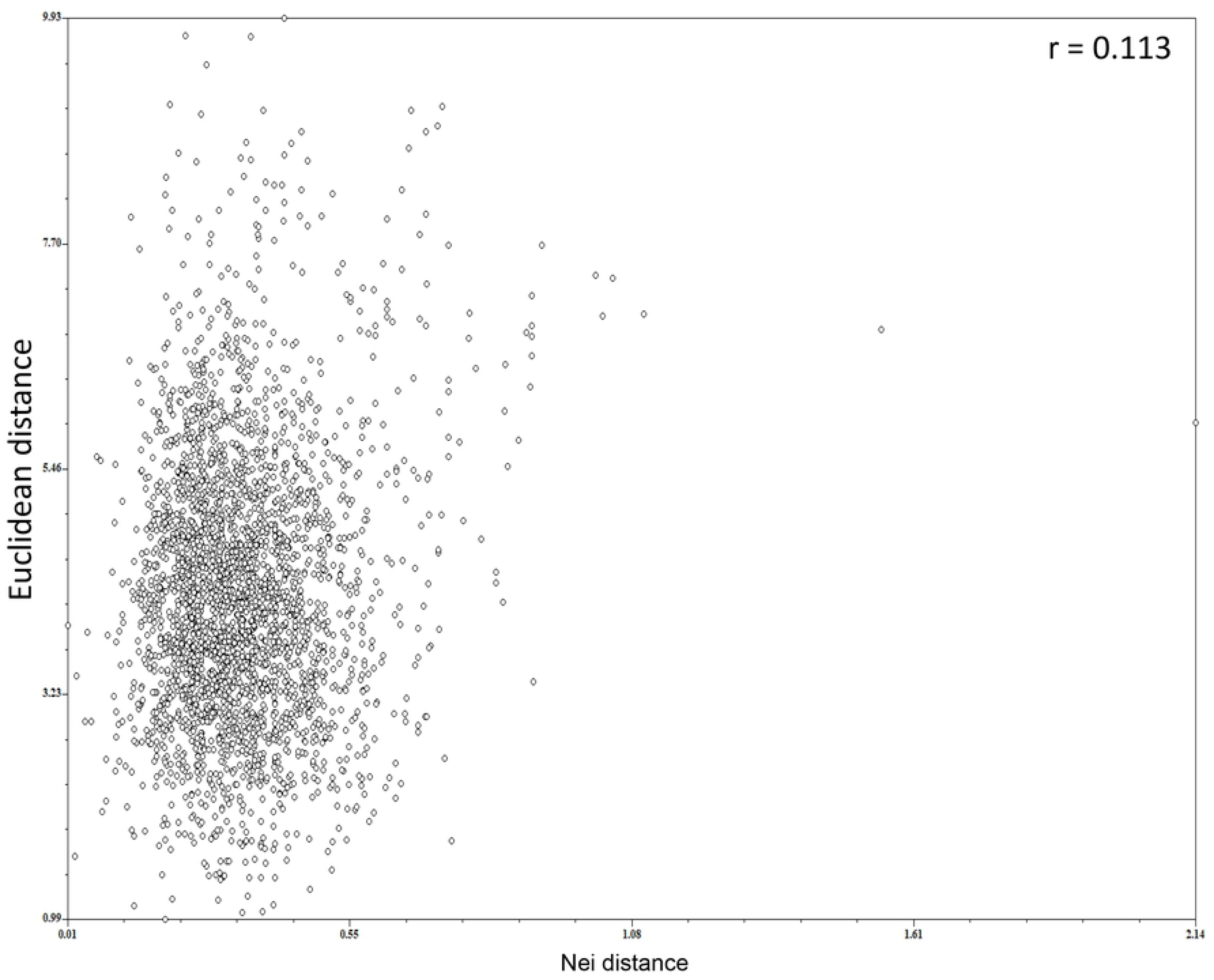
Pairwise relationship between Nei and Euclidean distance (r = 0.113) for 76 genotypes used in the diversity study. Nei distance was constructed based on ISSR marker data while Euclidean distance was constructed based on quantitative trait data.

## Discussion

Despite the severity of the virulence character of rice bacterial blight disease and regardless of the fact that host plant resistance is the best sustainable strategy to counter this age-old threat to rice production, no resistant variety so far has been developed in the country. However, a successful breeding program designed to develop resistant genotypes is largely influenced by the gene pool variations in the parental genotypes [45]. On the other hand, exploring genetic diversity of a collection of germplasm could enrich gene pool through introducing new gene(s) [24,46]. So, considering the importance of enrich and diversify the gene pool we explored a part of our landrace germplasm collection through morphological and molecular approaches.

Pathogenicity test exposed variant bacterial blight disease reaction among studied germplasm which may be an indication of the presence of divergent genes/alleles in the germplasm collection. Nonetheless, vast majority of the germplasm were found moderately susceptible to highly susceptible with only a single highly resistant germplasm, which was a clear indication of high virulent status of the *Xoo* strains under the current study [14–17,47].

The STS markers utilized in the study were found highly efficient to identify the resistant genes. Dominant resistant gene *Xa4*, first reported by [48], functioned via cell wall-associated kinase synthesis and confers a race-specific but plant growth stage nonspecific resistance towards *Xoo* strains [49-51]. Moreover, [51] deciphered the role of *Xa4* gene in improving certain quantitative traits such as, plant height. This may shed light on the reason behind the wide utilization on the *Xa4* gene in the rice breeding programs. In the current study, we identified around 55.41% germplasm (41 out of 74) harboring *Xa4* gene. Similarly, high presence rate of *Xa4* gene among the germplasm were also reported by [12,15,52,53]. Subsequently, among the two recessive *R* genes (*xa5* and *xa13*) under the current query, *xa5* was first identified by [48] and typically found in the Aus-Boro varieties of Bangladesh [11]. Recessive *R* gene *xa5* confers a race-specific resistance against *Xoo* by encoding a *TFII-λ* subunit [54]. However, in our study, around 20.27% germplasm (15 out of 74) showed to have recessive *xa5* gene. Similar to our finding, 34.29% germplasm were found to carry *xa5* gene in India [14], 17.5% in Pakistan [53] and 29.62% in Nepal [16]. Alternatively, [15,17] did not find of any *xa5* gene in Thai or Malaysian germplasm, respectively. These results may indicate the predomination of *xa5* gene in the Indian subcontinent. Nonetheless, another recessive *R* gene *xa13* was first reported in a landrace (BJ1) derived from Indian subcontinent [55]. Similar to *Xa4* and *xa5, xa13* is also a race specific *R* gene [56]. Both of *xa13* (resistant) and *Xa13* (Susceptible) allele encodes clade-III SWEET (Sugar will eventually be exported transporter) protein which is responsible for sugar efflux into the apoplast region of plant from cytoplasm [57-59]. Mutation in the -69 to -86 position of *OsSWEET11* or *Xa13* plays the central role in recessive *xa13* mediated resistance to *Xoo* [60]. However, in our study we identified that about 44.59% germplasm (33 out of 74) possessed resistant *xa13* allele. Regrettably, in some previous researches led by [14,15,17] were not able to detect any *xa13* gene in the germplasm derived from India, Thailand and Malaysia, respectively.

The first cloned *R* gene *Xa21* was categorized as an LRR-RKL (Leucine-rich repeat receptor-like kinases) gene [61]. Although, *Xa21* showed broad-spectrum resistance against *Xoo*, its activity largely depends on the plant growth stage [62-65]. However, *Xa21* had been identified as an effective and durable *R* gene against *Xoo* strains in Bangladesh [21,66], India [67,68], Vietnam [69], as well as in Korea [70]. Despite of its high effectiveness, *Xa21* appeared to be a rare gene in terms of presence in the germplasm [12,14,16,17,53]. In our study we found only one germplasm carrying *Xa21* gene. Interestingly, germplasm G3 carrying *Xa21* gene was the only germplasm after pyramided line IRBB60 with a highly resistant (HR) response against all the *Xoo* strains. This observation again proved the high effectiveness of *Xa21* gene against diverse *Xoo* strains. On the other hand, newly cloned executor *R* gene *Xa23* was identified from *Oryza rufipogon* by [71] and has been well recognized for it’s a broad-spectrum resistance against *Xoo* by inducing hypersensitive response in rice at all growth stages [72,73]. The seven base pairs mutation in the promoter site is the secret behind its activation by a conserved TALE (transcription activator-like effector) element, AvrXa23 encoded by *Xoo* strains [73,74]. Notably, in our study we identified 19 germplasm (25.68%) which carries resistant *Xa23* allele while, [12] did not find any resistant *Xa23* gene in Chinese cultivar.

Dominant but un-cloned *R* gene *Xa7* was first identified in DV85 rice cultivar and was reported to be predominate in Aus cultivars of Bangladesh [18,75,76]. Though it is not cloned yet, its durable resistance and effectiveness in high temperature makes it a desirable target gene in modern breeding programs especially in tropical country like Bangladesh [49,77]. In the present study, we identified maximum 83.78% germplasm (62 out of 74) which carries *Xa7* gene. These findings again indicated the predominance of *Xa7* gene in Bangladeshi germplasm. In contrast, [76] reported of 59.50% germplasm carrying *Xa7* gene in Pakistan, [15] reported of 11.61% germplasm carrying *Xa7* gene in Thailand, [16] reported of 23.33% germplasm carrying *Xa7* gene in Nepal and [12] reported only 4.91% germplasm carrying *Xa7* gene in China, in their respective study.

It is widely accepted that a single resistant gene, despite of its resistance range, will eventually lose its durability as pathogens are more prone to mutations [18,78]. On the other hand, rice varieties carrying multiple resistance genes provide a more effective and durable resistance towards diverse *Xoo* strains and hence are best suited for long term and large-scale cultivation. For example, [79] reported about *xa5*, *xa13* and *Xa21* gene combination providing superior resistance in Eastern India, [80] informed about *Xa4*, *Xa7* and *Xa21* gene combination showing higher resistance in Vietnam, [81] identified *xa5*, *Xa21*; *xa33*, *xa5*, *xa33*; *xa5*, *Xa21*; *Xa21*, *xa33* gene combinations conferring wider and higher resistance in Myanmar and Thailand, [5] found that *Xa4*, *xa5* and *Xa21* pyramided line exert a durable and long term resistance than a single *R* gene in Korea while, [36] reported about *Xa4* and *Xa7* gene combination was durable against *Xoo* at high temperature and [82] conveyed that *Xa4*, *xa5* and *Xa21* gene combination was superior than any of the single *R* gene. Likewise, in the present study we also observed that germplasm with multiple genes typically confer higher resistance than germplasm with single gene. Notably, among the different gene combinations recorded in our study, *Xa4*, *Xa7*, *xa13* and *Xa21* gene combination was most effective. We believe, the germplasm with diverse gene combinations, especially germplasm G3 (*Xa4*, *Xa7*, *xa13* and *Xa21*) and G43 (*Xa4*, *xa5*, *xa13* and *Xa23*), will play the role as harbinger in our future fight against *Xoo*.

Considering the importance of diverse collection of rice germplasm in future breeding programs, determination of genetic divergence among the germplasm is indispensable. Moreover, genetic divergence revealed through morpho-molecular markers paves the way of the breeders in sustainable management of biotic stresses like, *Xoo*. The average PIC and RP value of the ISSR markers observed in our study was moderate and analogous to the findings of [30,31,83]. The UPGMA dendrogram and two-dimensional scattered plot grouped the genotypes into 14 and 16 clusters, respectively. Both analyses showed almost similar clustering pattern and both were able to distinguish IRBB60 (G76), non-native to Bangladesh, from the rest of the germplasm. This was a clear indication of the effectiveness of the ISSR markers in resolving exotic germplasm collection. Moreover, genetic similarity matrix revealed that most of the germplasm had low genetic similarity with IRBB60, which further supported the effectiveness of the ISSR markers used in the experiment. Nonetheless, as the bacterial blight susceptible elite variety BRRI dhan28 had comparatively high genetic distance with germplasm G6 (carry *Xa4*, *Xa7*, *xa13*, *Xa23* genes), G17 (carry *Xa7*, *xa13* genes) and G20 (carry *Xa4*, *Xa7*, *xa13* genes). Hybridization could be made between BRRI dhan28 and these three germplasms to develop bacterial blight resistant variety [23,34,84].

Variations in quantitative traits among the bacterial blight resistant genotypes were highly significant (*p*≤0.01) as revealed by ANOVA analysis. This was an indication of the existence of wide inherited genetic variations among the germplasm. The UPGMA dendrogram based on quantitative data clustered the germplasm into 8 different clusters. Notably, in respect to the cluster mean values of quantitative traits we deduced that germplasm of the cluster-III had superior quantitative traits such as the lowest days to flowering and days to maturity, moderate panicle length, decent rate of effective tiller formation, lowest unfilled spikelet panicle^-1^ and highest 1000 grain weight. On the other hand, low plant height is considered as one of the most desirable traits in modern breeding programs as it improves lodging resistance of the plants [85,86]. Considering this fact, lower plant height with higher tiller number hill^-1^ made the germplasm from cluster-VIII a good candidate for future breeding programs. Nonetheless, PCA and UPGMA clustering patterns were almost similar in the both methods with a few exception [29,86,87].

## Conclusion

Pathogenicity test of 792 germplasm identified 74 potential germplasm conferring highly resistant to moderately resistant disease reaction against three highly virulent *Xoo* strains, *BXo69*, *BXo87* and *BXo93*. Further, molecular evaluation reveled that, the germplasm harbored 1 to 4 *R* genes of various combinations. Whereas, germplasm G3 carrying *Xa4*, *Xa7*, *xa13* and *Xa21* gene was the only germplasm showing highly resistant disease reaction against all three *Xoo* strains. Another germplasm G42 harbored 4 *R* genes (*Xa4*, *xa5*, *xa13* and *Xa23*) among which *xa5*, *xa13* and *Xa23* were some of the most effective *R* genes against *Xoo* strains from Bangladesh. This two germplasm could be recommended for future breeding programs to develop bacterial blight resistant superior varieties. Nevertheless, quantitative traits evaluation of the 74 selected germplasm identified two prominent clusters namely, cluster-III and cluster-VIII with multiple desired traits. Germplasm from these two clusters could also be recommended for future breeding programs. On the other hand, based on genetic distance gained from ISSR marker data, hybridization between elite variety BRRI dhan28 and germplasm G6/G17/G20 could also be a potential recommendation. Last but not the least, the outcome of this study would enrich and diversify the rice gene pool and would be useful for the development of bacterial blight resistant durable varieties.

## Acknowledgements

The authors acknowledge Genetic resource and seed division (GRSD), Bangladesh Rice Research Institute (BRRI), Gazipur, Bangladesh and International Rice Research Institute (IRRI), Philippines for their cooperation in providing the seed samples.

## Funding

This work was funded by NATP-2, PBRG, BLB sub-project (ID-091).

## Supporting information

**S1 Table.** List of 794 genotype used for pathogenicity study, their lesion length against three *Xoo* strains, disease reaction score and reaction result.

**S2 Table.** Disease reaction score, resistance status and STS marker amplification results of 74 potential germplasm.

**S3 Table.** Genetic similarity matrix based on Jaccard’s method showing pairwise genetic similarity among the 76 genotypes.

**S4 Table.** Different quantitative traits data of 76 genotypes.

## Notes

### Competing Interest Statement

The authors have declared no competing interest.

